# Real-time in vivo thoracic spinal glutamate sensing reveals spinal hyperactivity during myocardial ischemia

**DOI:** 10.1101/2023.03.11.531911

**Authors:** Siamak Salavatian, Elaine Marie Robbins, Yuki Kuwabara, Elisa Castagnola, Xinyan Tracy Cui, Aman Mahajan

## Abstract

Myocardial ischemia-reperfusion (IR) can cause ventricular arrhythmias and sudden cardiac death via sympathoexcitation. The spinal cord neural network is crucial in triggering these arrhythmias and evaluating its neurotransmitter activity during IR is critical for understanding ventricular excitability control. To assess the real-time *in vivo* spinal neural activity in a large animal model, we developed a flexible glutamate-sensing multielectrode array. To record the glutamate signaling during IR injury, we inserted the probe into the dorsal horn of the thoracic spinal cord at the T2-T3 where neural signals generated by the cardiac sensory neurons are processed and provide sympathoexcitatory feedback to the heart. Using the glutamate sensing probe, we found that the spinal neural network was excited during IR, especially after 15 mins, and remained elevated during reperfusion. Higher glutamate signaling was correlated with the reduction in the cardiac myocyte activation recovery interval, showing higher sympathoexcitation, as well as dispersion of the repolarization which is a marker for increased risk of arrhythmias. This study illustrates a new technique for measuring the spinal glutamate at different spinal cord levels as a surrogate for the spinal neural network activity during cardiac interventions that engage the cardio-spinal neural pathway.

**Graphical abstract:** 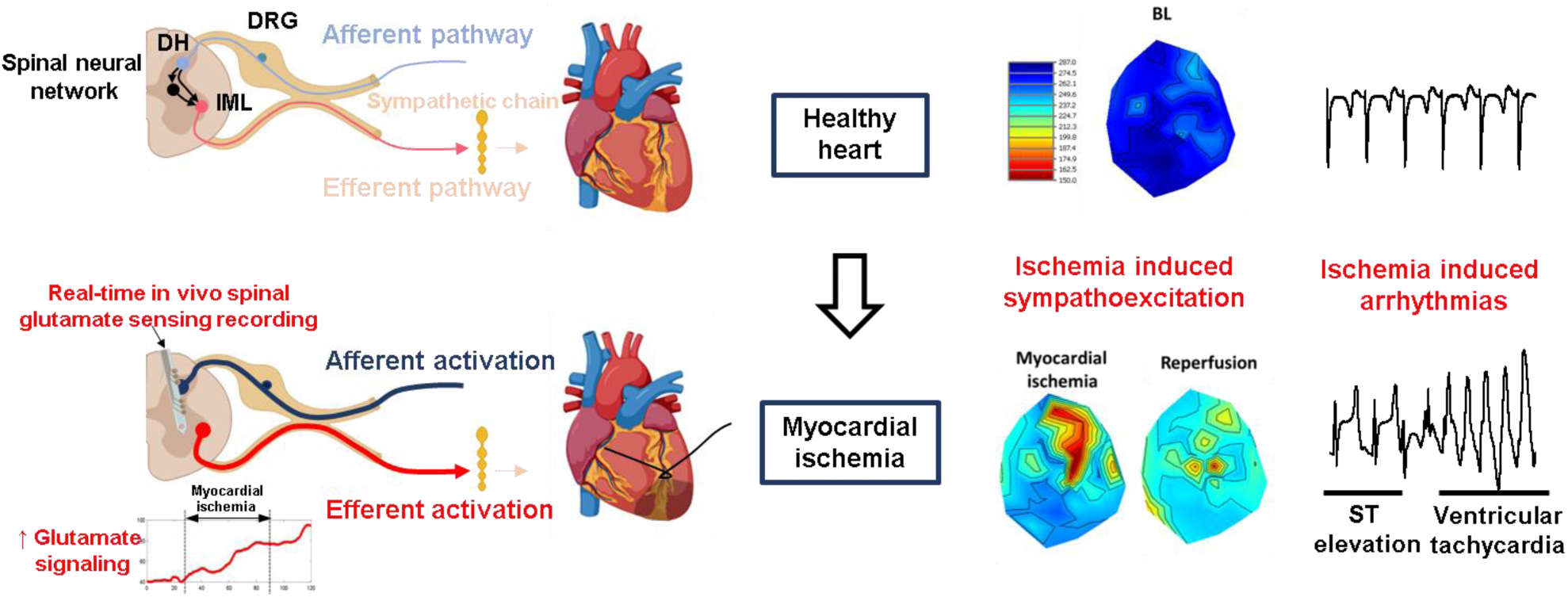

## Introduction

Myocardial ischemia-induced ventricular arrhythmias are the leading cause of sudden cardiac death (1). The cardiac autonomic nervous system (CANS) plays a key role in regulating cardiac function after myocardial ischemia(2). The autonomic balance between the sympathetic tone and parasympathetic tone when the heart is not stressed is the critical component of cardiac function regulation. When the heart is stressed, for instance during myocardial ischemia-reperfusion (IR) injury, the cardiac sensory neurons in the dorsal root ganglion transmit excitatory signals to the spinal cord. These excitatory signals are processed in the dorsal horn of the spinal cord and are translated to the activation of the sympathetic preganglionic neurons in the intermediolateral nucleus which results in the sympathoexcitation (3, 4). This sympathoexcitation is a necessary mechanism for the CANS to regulate the function of the heart during stress; however, if the cardiac stress persists, the excessive sympathoexcitation will cause autonomic imbalance that results in cardiac dysfunction. Autonomic imbalance is one of the main causes for the initiation and maintenance of the fatal arrhythmias (5–10).

It has been shown that short myocardial ischemia causes excitatory sensory response (11). However, it is not fully known how the cardiac sensory neurons respond to the longer myocardial ischemia (1 hr) and how the spinal cord neural network processes the cardiac sensory information. Myocardial ischemia has a different mechanism of action during different phases of ischemia which results in different sensory inputs to the spinal cord and hence different spinal cord network processing. Regional ischemia causes two phases of ventricular arrhythmia: i) Early phase which occurs between 5-7 mins after the coronary artery occlusion and results in mild intracellular and extracellular acidification and membrane repolarization, ii) the second phase happens between the 20-30 mins after the onset of myocardial ischemia while the ischemia-induced K^+^ and pH changes are stable (12). It is therefore important to understand how the spinal cord is processing the cardiac sensory information at different levels of the spinal cord and at different time points during myocardial ischemia.

Glutamate serves as the major excitatory neurotransmitter that is released in the spinal cord (13, 14) and its concentration can be used as a biomarker for the neural network activation level in the pathological state (15) but not the normal state due to the glutamate uptake mechanism (16). Some techniques have been tried to measure glutamate concentration in the spinal cord; for example, microdialysis was used to analyze glutamate concentrations in the pig spinal cord following aortic cross-clamping (17). However, a minimum amount of sample is needed for offline assay or HPLC analysis, which takes several minutes to be collected – in this case, one sample every 10 minutes (18). The resulting poor time resolution means that microdialysis does not provide time resolution sufficient to track the glutamate concentration change during IR events. Electrochemical glutamate detection with amperometry allows for sub-second time resolution. The current state-of-the-art glutamate electrochemical sensor is a ceramic microelectrode array consisting of a glutamate-sensitive electrode and a sentinel control electrode (19). We advance this technology by fabricating microelectrode arrays on a flexible SU-8 substrate for superior tissue integration (20). We also demonstrate decreased noise from movement artifacts with the flexible substrate, which is particularly important in a large preclinical animal model. Additionally, probes were fabricated with multiple glutamate sensing channels for simultaneous measurement across different laminae of the spinal cord. This is an important capability of multichannel sensing electrodes as different spinal cord laminae and nuclei process different information and have different functions (processing sensory information, motor neurons that innervate skeletal muscle for movement, preganglionic sympathetic neurons that innervate the paravertebral sympathetic chain to increase the sympathetic tone)(21). Further, the therapeutic approach of spinal cord stimulation during myocardial ischemia-reperfusion injury has differential effects on spinal neurons in superficial lamina vs. deeper lamina (22). To study the impact of cardiac IR on the lamina-specific neural networks in the spinal cord or for potential optimization of neuromodulation therapies, it is undeniably essential to investigate the spinal neurotransmitter activity using multichannel sensing probes.

In this study, we have evaluated the spinal neural network activity during IR injury using a newly developed custom-made flexible probe that is capable of measuring glutamate concentrations at different levels of the spinal cord. The glutamate detection of the developed probe is almost instantaneous and can measure the glutamate concentration continuously. Using this developed continuous spinal glutamate measurements, we have shown that, the glutamate concentration increases during the myocardial ischemia and reperfusion injury which causes sympathoexcitation and increases the risk of ventricular arrhythmias. A large animal porcine model was used in this study due to its high translatability to humans in terms of the cardiovascular system and the CANS.

## Results

### Design of the flexible microelectrode array

The final microelectrode array (MEA) design is schematically represented in Figure 1A. The flexible MEAs were successfully fabricated on the SU-8 flexible substrate following a direct-writing maskless photolithography process (Figure 1B). SU-8 has demonstrated excellent chronic performance as a neural probe substrate(23–27). The device is composed of a singular shank (140 µm wide and ∼16 µm thick) with 6x 100 µm diameter platinum electrode sites, organized into two groups of three electrodes each. The total length of the shank is 6.5 mm to easily target the spinal cord dorsal horn and IML of the pig. An anchor hole was patterned at the shank tip for the insertion of a 50 µm sharpened tungsten shuttle, which facilitates the handling and implantation of the flexible device into the spinal cord (Figure 3C). The flexible MEAs are connected to the PCB using a zero-insertion force (ZIF) connector (Figure 3A), to be interfaced with characterization and recording systems. The PCB is sealed with epoxy to protect the connection from fluids and blood exposure. The final MEA is highly flexible and can be bent significantly without breaking (Figure 3A).

**Figure 1.**
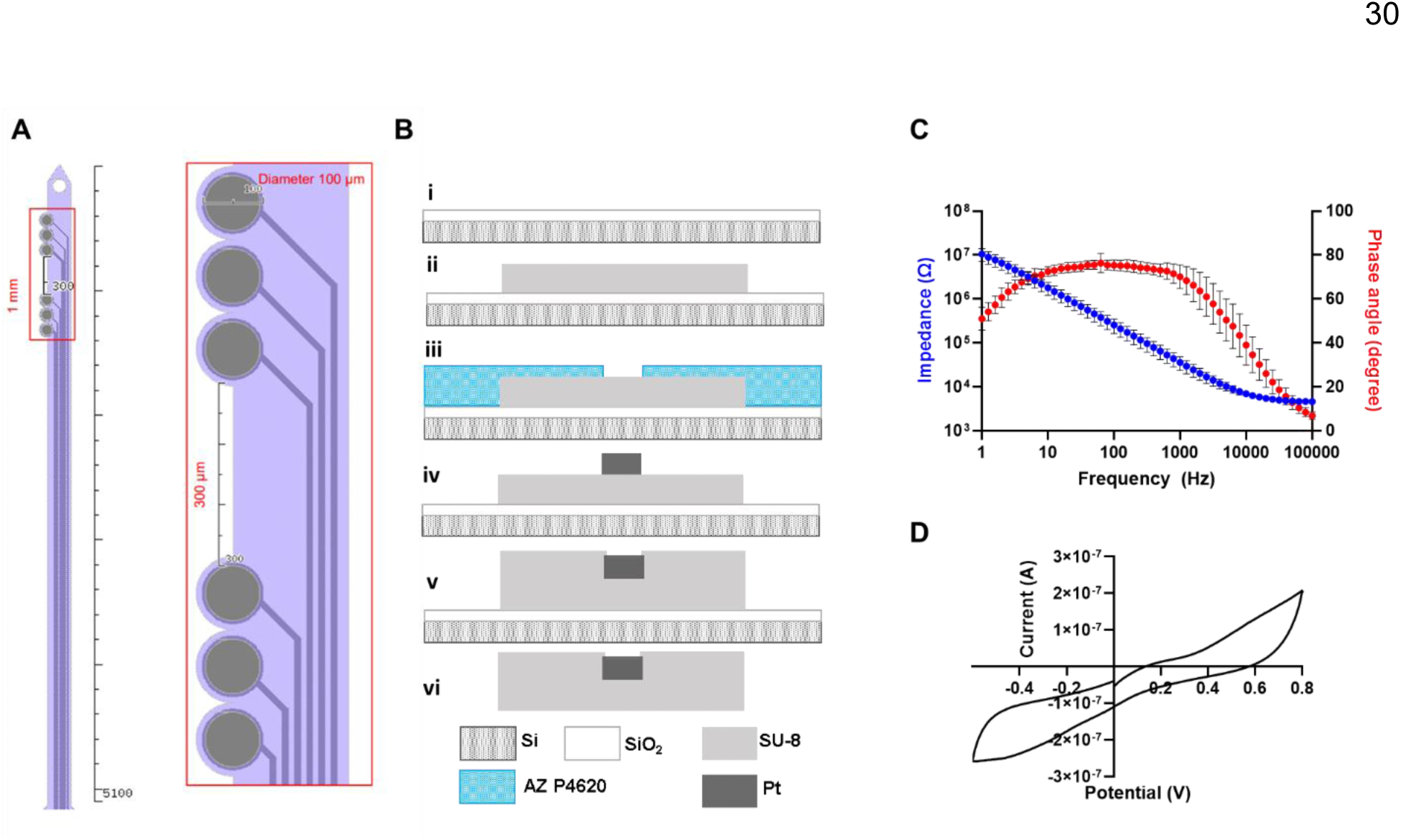
Electrodes design. **(A) Schematic of the probe design**. (B) Graphic of the photolithography process. (C) Electrochemical impedance spectroscopy (EIS) and phase angle of all 6 sites on 3 probes with high reproducibility within electrodes and between electrodes. (D) CV of an electrode

Electrochemical impedance spectroscopy (EIS) of all 6 electrode sites on three arrays (for a total of 18 sites) show excellent reproducibility both between sites on a single MEA as well as across multiple MEAs (Figure 1C). The average impedance at 1 kHz of these sites was 36 ± 3 kΩ (mean ± SEM). Similarly, cyclic voltammetry measurements between -0.6 and 0.8 V vs. Ag/AgCl, measured at 0.1 V/s have an average charge storage capacity of 3.5 ± 0.3 µC (Figure 1D) *Sensitivity of the glutamate electrodes*

The MEAs were modified to be glutamate-sensitive electrodes. A schematic of the layers on the electrode sites is shown in Figure 2A. The final MEA contains electrode sites that are either sensitive to glutamate (the GluOx sites) or are controls (the sentinel sites). In a GluOx site, GluOx bound to the surface of the electrode reacts with glutamate to produce H_2_O_2_, which is oxidized at the Pt electrode surface to produce a current. The sentinel sites are not sensitive to glutamate but will react to ambient H_2_O_2_ in the tissue, along with any other potential interferences. The final glutamate-derived signal is calculated as the difference between the GluOx sites and the sentinel site. The two types of sites are constructed identically, except the sentinel sites do not have GluOx crosslinked on their surface. Full details of sensor construction are in the Methods; to briefly summarize the process, a size-exclusion screening layer of m-phenylenediamine (mPD) was first electrochemically deposited onto the surface. Subsequently, either a glutamate oxidase (GluOx)-containing layer, or a control sentinel layer was carefully drop casted under a microscope onto each electrode site and allowed to crosslink overnight.

**Figure 2.**
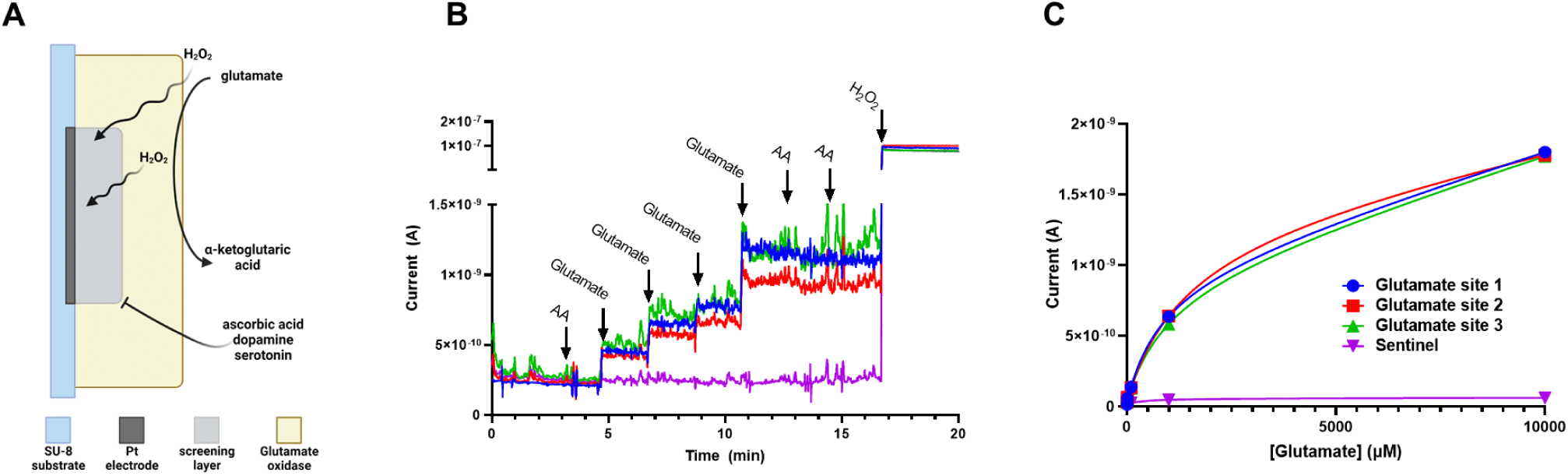
Glutamate sensor performance. (A) Schematic of how the glutamate sensors work. (B) Sensors in a stirred solution of PBS. Electrodes don’t respond to additions of ascorbic acid. The glutamate sites respond to glutamate but not to the addition of ascorbic acid, while the sentinel electrode does not respond to both glutamate and ascorbic acid. All sites respond to hydrogen peroxide. (C) Calibration curve shows a wide range of activity for the glutamate sites and no activity for the sentinel.

**Figure 3.**
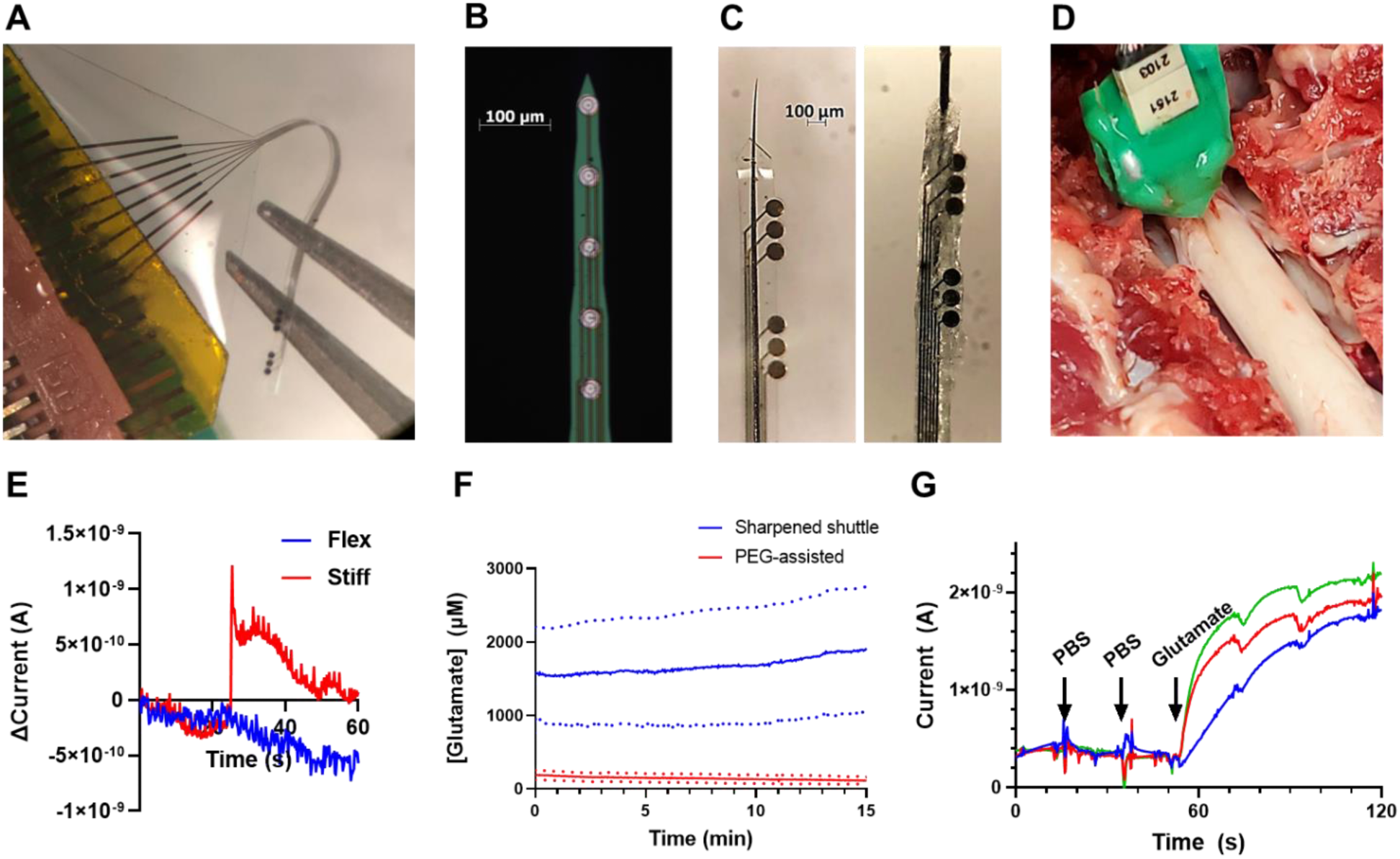
Performance of flexible probe. (A) Photo demonstrating the flexibility of the probe. (B) A photo of the stiff Neuronexus probe. (C) Left: Tip of an electrode assembled with the sharpened tungsten. Right: polyethylene glycol (PEG) assisted probe (D) A photo of the electrode inserted into the pig spinal cord. (E) Data were recorded during motion artifacts using a stiff acute Neuronexus probe and our flex probe, both inserted and recorded at the same time. (F) The sharpened shuttle method results in significantly more detected glutamate in vivo due to the PEG covering the electrodes’ surface. (G) Ex vivo injections of PBS and glutamate into a spinal cord. The electrodes do not sense the PBS but the injected glutamate is sensed by the glutamate sensor probe.

The final electrodes were highly selective to their desired target. Both the GluOx and sentinel sites have no response to the main interfering compound, ascorbic acid. However, the GluOx sites (red, green, and blue traces) respond robustly to repeated addition of glutamate to the solution and as expected, the sentinel site (purple) does not respond (Figure 2B). However, as expected, every site responds to H_2_O_2_. When calibrated, the GluOx sites show sensitivity to glutamate across a wide range of concentrations between 10 µM and 1 mM (blue, green, and red traces), while the sentinel site shows no response (purple trace) (Fig 2C). All final in vivo concentrations presented here were calculated using a post-calibration curve constructed by exposing the electrodes to standard concentrations of glutamate after the electrode was removed from the spinal cord.

To validate the sensing capabilities in tissue, we inserted the electrodes into a piece of pig spinal cord that had been carefully dissected from the animal. A needle was inserted near the electrode, and a series of injections of vehicle phosphate-buffered saline (PBS) or glutamate were performed. As expected, the electrodes did not respond to the vehicle; however, there was a robust response to the injection of glutamate (Figure 3G).

We have also tested the sensing capabilities of our probe in a sham condition and observed a slight decrease in the detected current over 2 hours of recordings (Supplemental Figure 2A). The sham recording was performed after a stabilization period of at least 30 minutes. We attribute this slow, linear decrease in current to the decrease in sensitivity of our electrode over time, which could be the result of enzyme degradation or biofouling. We have corrected this by fitting a line to the first 30 minutes of the baseline data and then subtracting the slope of the fitted line to correct the data for the slight decrease in sensitivity over time in vivo (Supplemental Figures 2A and 2B). Supplemental Figure 2C shows the glutamate concentration changes during the 30-minute baseline after the correction of the loss of glutamate sensitivity.

### Performance of the flexible probe during movement

Because ambient movement due to breathing, heartbeat, etc. is increased in a large animal model, we fabricated the MEA on a highly flexible SU-8 substrate. Flexible probes have been demonstrated to reduce inflammatory tissue responses caused by micromotion of the brain following implantation in rodents (20). However, in a large animal model, continuous passive movements caused by a biological process such as breathing are much more dramatic. The spinal cord itself also presents an additional challenge; touching the spinal cord can inadvertently stimulate neuron firing resulting in muscle spasms. These spasms are common during electrode insertion and may damage the tissue and break silicon-based probes that are brittle and fragile, like those we have used previously to study the spinal cord (3, 4). Our previous experience with stiff probes led us to believe that a flexible probe will be more likely to better accommodate the animal’s movement.

To investigate this, we inserted both a flexible and a stiff probe into the spinal cord. The stiff probe was a commercially available probe purchased from Neuronexus with 30 µm platinum electrode sites, while the flex probe was a specially fabricated MEA that also had 30 µm platinum electrode sites (compared to the 100 µm sites used in the rest of this work) for an accurate comparison to the stiff probe. Sites from both electrodes were connected to a multichannel potentiostat for simultaneous amperometric measurements at 0.7 V vs. Ag/AgCl reference. After a 30-minute stabilization time, the spinal cord was lightly brushed with a cotton swab to induce a mild muscle spasm. This movement induced a sudden and dramatic change in the baseline of the stiff probe (red trace) but not in the baseline of the flex probe (blue trace) (Figure 3E) This test was repeated multiple times with a five-minute re-stabilization period between each trial (Supplemental Figure 1B). The response of the stiff probe decreased with each subsequent spasm, presumably due to damage and loss of tissue cohesion around the stiff probe.

### Optimization of the spinal cord insertion procedure

While flexible probes can move with the tissue during micromotions and improve device-tissue integration(20, 28–30), insertion of these flexible devices into the tissue becomes more difficult(31–33)

. Flexible materials often are too soft to penetrate tissue on their own, so shuttle systems are common to aid insertion (20, 27, 34, 35).

To that end, we compared two insertion methods from the literature to determine the optimal procedure for neurochemical sensing in the spinal cord. In one scheme, a 50 µm diameter tungsten guide wire was threaded through a 100 µm insertion guide hole at the tip of the SU-8 shank. The wire was then glued to the MEA with 30% polyethylene glycol (PEG) (Figure 3C). In the second procedure, the hole in the SU-8 shank was 40 µm wide, and a 50 µm SU-8 wire was sharpened by applying 5 V vs. Ag/AgCl and repeatedly dipping it into a solution of 5 M potassium hydroxide to etch the surface. The final needle-like wire was threaded through the insertion hole without the PEG glue (Figure 3C).

When comparing the PEG-assisted and tungsten shuttle insertion methods, the PEG-assisted insertion technique (red, n=8 electrode sites) has a signal several orders of magnitude lower than the sharpened-shuttle method (blue, n=4 electrode sites) (Figure 3F). Figure S1A shows the mean ± SEM of all functional, glutamate-detecting electrodes (n=12 GluOx sites in n=6 pigs).

### Spinal cord hyperactivity during myocardial ischemia

Glutamate signaling was measured as a biomarker for spinal cord neural activity. We measured the glutamate concentration during the 30 minutes of baseline recording (in 5 mins intervals) and we did not see a difference in glutamate concentration during these 30 minutes (n=12 electrodes, P= 0.61, Friedman test) (Supplemental Figure 2C). Glutamate percentage change from baseline was assessed during the baseline, every 15 minutes of ischemia, and the 15 minutes of reperfusion. The percentage change in glutamate was not significant during the first 15 minutes of the LAD ischemia (BL: -0.03 [-1.85, 1.90] % to LAD15: 11.04 [4.75, 17.94] %, P= 0.63) Glutamate concentration started to increase significantly between the 15-30 minutes of ischemia (BL: -0.03 [-1.85, 1.90] % to LAD30: 24.40 [12.81, 259.77] %, P= 0.03) (Figure 5A, 5C). Glutamate augmentation persisted during the rest of the myocardial ischemia during 30-45 minutes (BL: -0.03 [-1.85, 1.90] % to LAD45: 53.788 [19.46, 285.46] %, P= 0.0002) and 45-60 minutes of LAD (BL: -0.03 [-1.85, 1.90] % to LAD60: 69.64 [29.75, 205.63] %, P< 0.0001). Glutamate concentration did not decrease during the 15 minutes of the reperfusion and was still elevated compared to the baseline (BL: -0.03 [-1.85, 1.90] % to Reperfusion: 69.35 [36.87, 159.02] %, P <0.0001) (Figure 5A, 5C).

### Shortening of the activation recovery interval during IR injury

The activation recovery interval (ARI) is a surrogate for the action potential duration (Figure 4D). The action potential duration becomes shorter when the sympathetic tone is high and therefore the ARIs shortens during the sympathoexcitation. In this study, the ARIs were shortened (285.06 ± 15.75 ms to 256.08 ± 13.88 ms, P=0.01) during myocardial ischemia (Figure 6B) which shows the augmentation in the sympathetic tone. The ARI shortening occurred within the first 5 minutes of the ischemia and persists until the termination of the myocardial ischemia (Figure 5B, 5C). The sympathetic activation was not mitigated during the reperfusion as the ARI was still shorter during the reperfusion compared to the baseline (285.06 ± 15.75 ms to 260.29 ± 19.01 ms, P=0.04) (Figure 6B).

**Figure 4.**
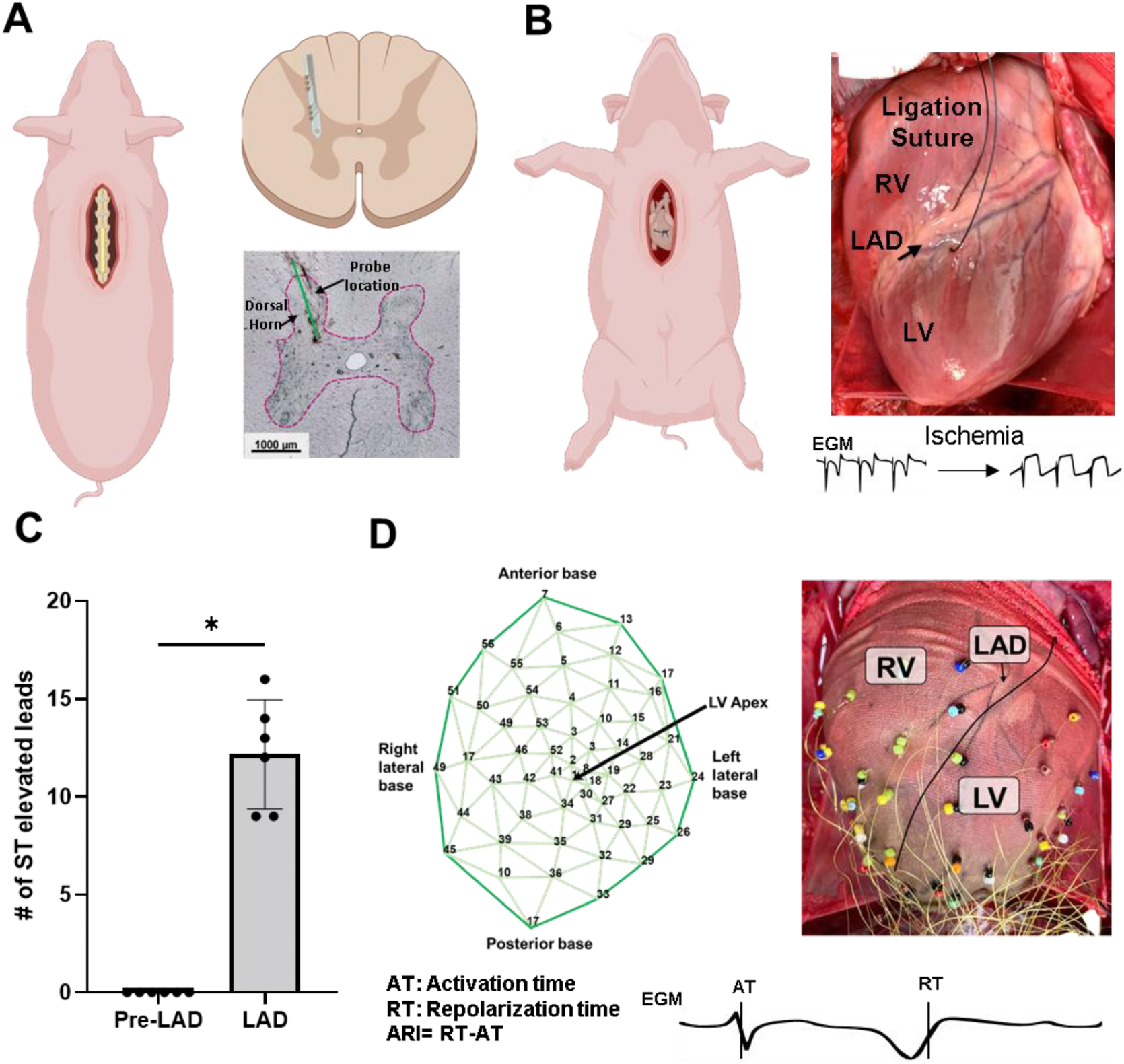
Simultaneous in-vivo spinal glutamate and cardiac electrophysiological recordings. (A) Following T1-T4 laminectomy, the glutamate sensing probe was inserted in the dorsal horn region of the spinal cord at the T2-T3 level. (B) Myocardial ischemia was induced by ligating the suture that was placed around the left anterior descending (LAD) coronary artery. Myocardial ischemia was confirmed by observing the ST-elevated electrograms (EGM) in the ischemic region. LV/RV: left/right ventricle (C) Number of ST elevated leads increased during myocardial ischemia (n =6 animals Pre-LAD vs. 6 animals during LAD, *P* = 0.031, Wilcoxon test) (D) Following sternotomy, the 56-channel sock electrode was placed around the heart to record the electrocardiogram. The activation recovery interval (ARI) was measured by subtracting the activation time (AT) from the recovery time (RT).

**Figure 5.**
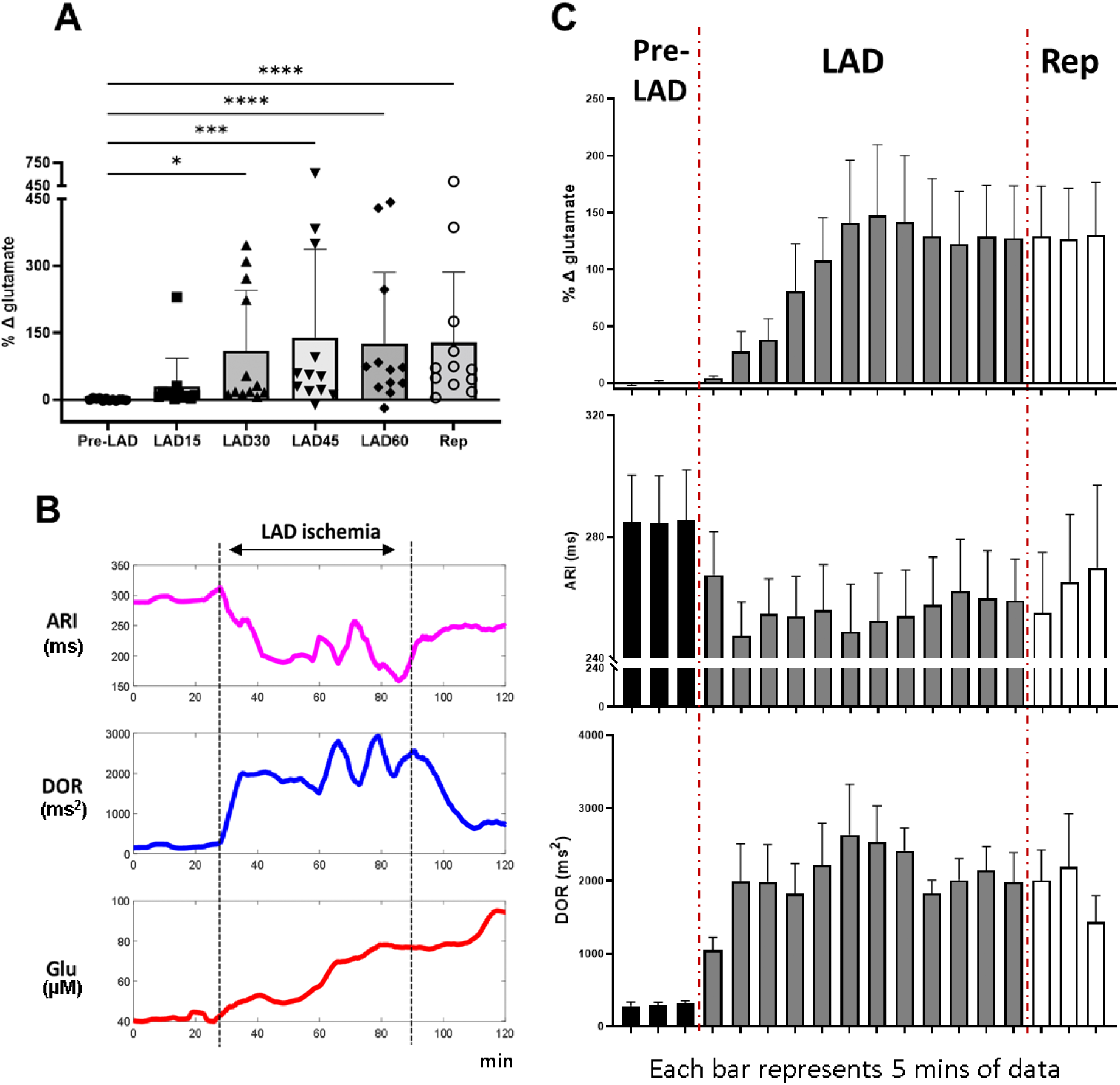
Simultaneous in-vivo spinal glutamate and cardiac electrophysiological recordings. (A) Myocardial ischemia-reperfusion injury caused an augmentation in the percentage change of glutamate from baseline (Friedman test, P <0.0001). Glutamate did not change during the first 15 minutes of the left anterior descending (LAD) coronary artery occlusion (n = 12 electrodes Pre-LAD vs. 12 electrodes during LAD15, *P* = 0.63, Dunn’s multiple comparisons test). LAD between 15-30 mins (n = 12 electrodes Pre-LAD vs. 12 electrodes during LAD15, *P* = 0.032, Dunn’s multiple comparisons test), LAD between 30-45 mins (n = 12 electrodes Pre-LAD vs. 12 electrodes during LAD15, *P* = 0.0002, Dunn’s multiple comparisons test), LAD between 45-60 mins (n = 12 electrodes Pre-LAD vs. 12 electrodes during LAD15, *P* <0.0001, Dunn’s multiple comparisons test) and reperfusion (n = 12 electrodes Pre-LAD vs. 12 electrodes during LAD15, *P* <0.0001, Dunn’s multiple comparisons test) caused an increase in the percentage change of glutamate from baseline. (B) Representative glutamate (Glu), activation recovery interval (ARI), and dispersion of repolarization (DOR) response during LAD ischemia. (C) Glutamate and DOR increased during the LAD ischemia and reperfusion, while the ARIs decreased during the ischemia-reperfusion injury.

**Figure 6.**
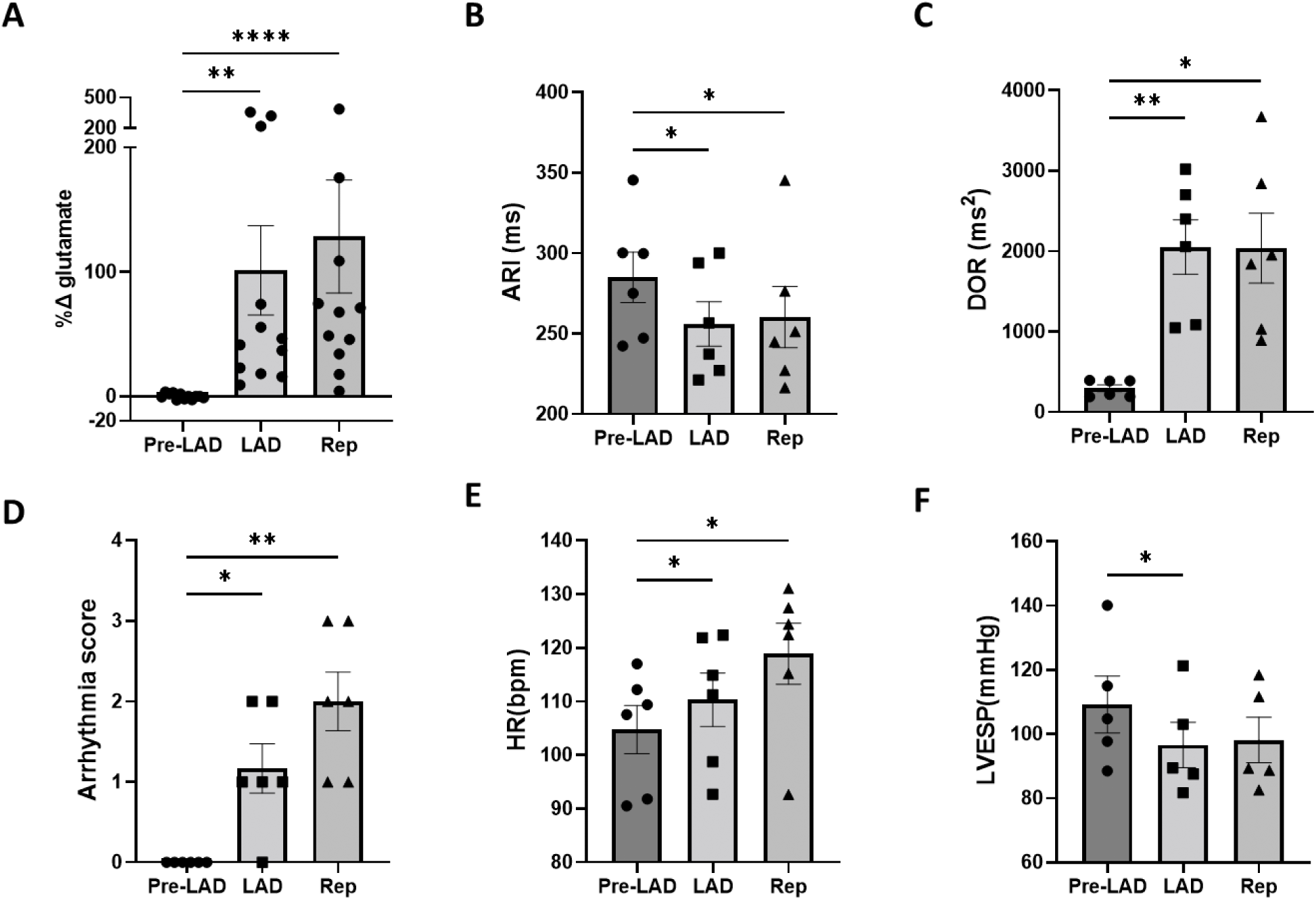
Physiological responses to the myocardial ischemia. (A) Left anterior descending (LAD) myocardial ischemia (n = 12 electrodes Pre-LAD vs. 12 electrodes during LAD, *P* =0.0085, Dunn’s multiple comparisons test) and reperfusion (Rep) (n = 12 electrodes Pre-LAD vs. 12 electrodes during reperfusion, *P* <0.0001, Dunn’s multiple comparisons test) caused an increase in the percent change of the glutamate concentration from baseline. (B) Activation recovery interval (ARI) was shortened during the LAD ischemia (n = 6 pig Pre-LAD vs. 6 pigs during LAD, *P* = 0.041, Dunn’s multiple comparisons test) and reperfusion (n = 6 pig Pre-LAD vs. 6 pigs during reperfusion, *P* = 0.0078, Dunn’s multiple comparisons test), demonstrating the sympathoexcitation. (C) Dispersion of repolarization (DOR) increased during the LAD ischemia (n = 6 pig Pre-LAD vs. 6 pigs during LAD, *P* = 0.0078, Dunn’s multiple comparisons test) and reperfusion (n = 6 pig Pre-LAD vs. 6 pigs during reperfusion, *P* = 0.041, Dunn’s multiple comparisons test), demonstrating the higher arrhythmogenicity. (D) Arrhythmia score increased during the LAD ischemia (n = 6 pig Pre-LAD vs. 6 pigs during LAD, *P* = 0.022, Dunnett’s multiple comparisons test) and reperfusion (n = 6 pig Pre-LAD vs. 6 pigs during reperfusion, *P* = 0.0049, Dunnett’s multiple comparisons test) as evidence of increased ventricular arrhythmias. (E) Heart rate was increased during the LAD ischemia (n = 6 pig Pre-LAD vs. 6 pigs during LAD, *P* = 0.033, Dunnett’s multiple comparisons test) and reperfusion (n = 6 pig Pre-LAD vs. 6 pigs during reperfusion, *P* = 0.038, Dunnett’s multiple comparisons test) and (F) Left ventricular end-systolic pressure (LVESP) were reduced during the myocardial ischemia (n = 5 pig Pre-LAD vs. 5 pigs during LAD, *P* = 0.0103, Dunnett’s multiple comparisons test), but not during the reperfusion (n = 6 pig Pre-LAD vs. 6 pigs during reperfusion, *P* = 0.303, Dunnett’s multiple comparisons test)

### Increased dispersion of repolarization during the myocardial ischemia

Dispersion of repolarization (DOR) is a potential marker for arrhythmogenicity. Increased DOR indicates an increased arrhythmia potential. The DOR started to increase within the first 5 minutes of the myocardial ischemia (Figure 5B, 5C) and stayed elevated by the end of myocardial ischemia (296.42 ± 42.18 ms^2^ to 2051.48 ± 337.91 ms^2^, P= 0.01) (Figure 6C). The DOR was not decreased during the 15 mins reperfusion (296.42 ± 42.18 ms^2^ to 2039.72 ± 435.00 ms^2^, P=0.02) and therefore the risk of arrhythmia events was still high during the reperfusion (Figure 6C).

### Increased arrhythmias during myocardial ischemia

Arrhythmia scores were measured during myocardial ischemia and reperfusion to evaluate the occurrence of premature ventricular contractions and ventricular arrhythmias. The arrhythmia score was augmented during ischemia (0 ± 0 to 1.17 ± 0.31, P=0.02) and reperfusion (0 ± 0 to 2.00 ± 0.37, P=0.005) (Figure 6D).

### Hemodynamic changes during myocardial ischemia

Heart rate and left ventricular end-systolic pressure were assessed during the myocardial ischemia and 15 mins reperfusion. Heart rate increased during the myocardial ischemia (104.76 ± 4.49 bpm to 110.31 ± 4.99 bpm, P=0.03) and reperfusion (104.76 ± 4.49 bpm to 118.88 ± 5.67 bpm, P= 0.04) (Figure 6E). Left ventricular end-systolic pressure decreased during the ischemia (109.23 ± 8.84 mmHg to 96.61 ± 7.08 mmHg, P=0.01). There was not a significant change in left ventricular end-systolic pressure from baseline to reperfusion (109.23 ± 8.84 mmHg to 98.15 ± 7.08 mmHg, P=0.30) (Figure 6F).

## Discussion

In this study, we have investigated the effect of myocardial ischemia on the neural processing at the T2-T3 level of the spinal cord by measuring the spinal dorsal horn glutamate concentration via amperometry technique. We have developed a new tool with a flexible probe to measure the spinal glutamate signaling in-vivo in a large animal model. Then we used this developed technique to measure the spinal glutamate signaling during myocardial ischemia and reperfusion which causes ventricular arrhythmias that are a leading cause of death. We have shown that the glutamate concentration increases during the myocardial ischemia, and it persists during the reperfusion and is associated with sympathoexcitation and increases in myocardial substrate excitability.

### Flexible glutamate-sensing probe

A flexible probe was developed in this study for minimizing probe insertion trauma to the spinal cord during the glutamate probe insertion. Several studies have evaluated the effect of probe stiffness on neuronal tissue injury, and they support the use of softer and more flexible probes (20, 28). Further studies confirmed the advantage of using flexible probes in terms of their capabilities to move with the neural tissue during micromotion that might be caused by cardiac pulsation or respiration (20, 28, 36, 37). In the current study, we have also shown that the flexible probe is resistant to movement artifacts during larger movements caused by other stimuli. This demonstrates the value of a flexible probe in large animal models, where motion caused by breathing, heartbeat, etc. is more intense than in a rodent model. Large tissue displacements around a stiff probe can cause damage to both the tissue and the electrode itself; a flexible probe can minimize these effects (38). Another study investigated the effect of probe stiffness in modulating the glial cells and their results indicate that glial cells respond greater to mechanical signals that are caused by stiff probes. (39). The flexible probe design provides a great opportunity for more reliable and robust recordings without causing significant trauma in the neural tissue. Though isolated use of flexible glutamate sensors has been reported as a proof-of-concept in rodents by a few investigators, we provide a comprehensive testing and in-vivo study of the sensors in a large pre-clinical disease model (40–44). The flexible glutamate sensors were successfully produced with the fabrication process described in Figure 1, followed by the coating procedure illustrated in Figure 2. The flexible sensors demonstrated selectivity to glutamate as well as sensitivity in the range of the concentration detected in vivo. To our knowledge, this is the first report of a flexible glutamate sensor used in a large animal model. An experiment to analyze the effect of physical motion on the electrode determined that the dramatic motion artifacts created by movement in a stiff probe were almost completely absent in a flexible probe (Figure 3E).

One of the major drawbacks of flexible sensors in vivo is the difficulty of inserting a probe that may buckle and fail to penetrate the neural tissue. Stiff shuttles or temporary stiffening approaches are often employed to aid in probe insertion into tissue and prevent buckling (20, 27, 29, 45, 46). Here, we compared two common techniques from the literature: a sharpened shuttle “needle-and-thread” approach, and PEG-assisted insertion, where PEG is used as a dissolvable glue to attach the flexible probe to the stiff shuttle. We found that the sharpened shuttle approach resulted in a much more robust detection of glutamate. We hypothesize this to be the result of two phenomena. First, adhering the flexible probe to the shuttle required a very thick layer of PEG that may not be effectively washed away for hours *in vivo*. The glutamate sensors require glutamate to be able to diffuse to and come into contact with the coating surface to react. While PEG is very soluble in an aqueous environment, residual PEG may still cover part of the sensing surface, which will decrease the effectiveness of the sensor. Second, the PEG-assisted insertion was much more difficult. In our hands, inserting the electrode with the PEG-assisted method frequently required several attempts. This may have resulted in both damage to the tissue as well as damage to the enzyme coatings on the sensor. The sharpened shuttle approach, therefore, resulted in more robust signals due to the ease of insertion and a more pristine electrode surface.

Our multielectrode array sensor was fabricated with 6 total channels – 3 glutamate sensing sites, and 3 redundant sentinel sites. By performing measurements with a 4-channel multi-potentiostat, we monitored the 3 glutamate-sensitive channels and one sentinel channel simultaneously. At different sites, we detected different concentrations of glutamate, possibly indicating local spatially-resolved concentration differences, and at some locations, we detected no glutamate at all despite the post-calibration confirming the sensor is functional (Supplemental Figures 2D and 2E). Future experiments to correlate the position of each electrode to the detected glutamate concentration will provide valuable insight into lamina-specific glutamatergic activity in the spinal cord.

### Sympathoexcitation and Myocardial Ischemia-induced Ventricular Arrhythmias

A large animal porcine myocardial ischemia-reperfusion model was used in this study. This model is well-established and studied by our group and others and it closely mimics the myocardial ischemia-reperfusion condition in humans. The myocardial ischemia was created and confirmed by at least 9 (out of 56) ST-segment elevated leads (Figure 4C). The ischemia-reperfusion injury increased the sympathoexcitation which was manifested by the shortening of the ARIs (4, 47). IR also increased the DOR which is associated with higher myocardial electrical heterogeneity and increased risk of arrhythmias (47–50). In this study, two pigs (out of 8) experienced ventricular tachyarrhythmias during myocardial ischemia and needed to be defibrillated. The occurrence of VT in these two pigs provides clear evidence of the risk of fatal ventricular arrhythmias during our large animal myocardial ischemia model. Furthermore, we have shown that the arrhythmia score has significantly increased during myocardial ischemia. The arrhythmia score is calculated based on the number of ventricular arrhythmias, including premature ventricular contractions and ventricular tachycardia or fibrillation episodes.

### Spinal hyperexcitation during myocardial ischemia-reperfusion injury

Mechanosensitive, chemosensitive, and multimodal afferent neurons transduce myocardial ischemia. These excitatory sensory signals are transmitted to the dorsal root ganglion and vagal nodose ganglia (11, 51). The dorsal root ganglia neurons transmit this afferent excitatory signal to the spinal dorsal horn neurons which will be processed in the spinal dorsal horn neurons and will activate the sympathetic preganglionic neurons in the IML region. Glutamate is a critical neurotransmitter for these spinal excitatory synaptic transmission (13, 14). Although the uptake mechanism makes it hard to detect the glutamate during the normal state (16), it is possible to measure the extracellular glutamate concentration during the hyperexcitation in a pathological state (15). In this study, we have simultaneously recorded the spinal glutamate signaling and cardiac electrophysiological signals in-vivo to provide evidence that the spinal glutamate signaling can be used as a biomarker for acute myocardial ischemia severity detection. We have shown that the glutamate concentration increases significantly between 15-30 mins of the myocardial ischemia and it stays elevated even during the reperfusion. The timing of the significant augmentation of the spinal glutamate correlates with the second phase of ventricular arrhythmias during myocardial ischemia between 20-30 mins after the onset of myocardial ischemia while the ischemia-induced K^+^ and pH changes are stable (12).

#### Limitation

Our study has some limitations that need to be addressed for clinical translation. First, neural activity can be suppressed in studies performed on anesthetized subjects. While we used alpha-chloralose as an anesthetic agent to minimize the effect of the anesthesia on the excitation of neurons, there might still be some suppression effects on the neural network. In this study, we have used acute implants for a proof-of-concept study. Future studies are needed to use this technique in a chronically implanted model and evaluate the efficacy of the probes in a chronically implanted model. We have shown that the glutamate increases significantly during the 15-30 minutes of ischemia.

## Methods

### Flexible Microelectrode Arrays Fabrication

A 4-in Si wafer with a 100 µm thick SiO_2_ layer (University Wafer Inc. Boston, MA, USA) was first cleaned with acetone, isopropanol, and DI water sequentially. The wafer was then dried with a N spray gun, heated on a hot plate at 150°C for 5 min, and treated with O_2_ plasma using a reactive ion etcher (RIE, Trion Phantom III LT) for 2 min at 300 mTorr pressure and 150 W power. The cleaned wafer was spin-coated with SU-8 2015 (MicroChemicals, Ulm, Germany) at 5000 rpm for 1 min and soft baked at 65°C for 3 min and 95°C for 5 min. Then, the wafer was exposed using a direct-writing maskless aligner (MLA, MLA100 Heidelberg Instruments) with a dose of 350 mJ/cm^2^ to define the bottom insulation layer. After exposure, the wafer was first post-baked at 65°C for 3 min and 95°C for 3 min, then developed using an SU-8 developer (MicroChemicals) for 1 min and cleaned with isopropanol and DI water. The patterned SU-8 was subsequently hard baked at 200°C, 180°C, and 150°C for 5 min each and allowed to cool down below 95°C.

After cleaning, the wafer was spin-coated with an AZ P4620 photoresist (MicroChemicals) at 5300 rpm for 1 min and baked at 105°C for 5 min, as previously described (1). After soft baking, the wafer was exposed using MLA with a dose of 700 mJ/cm^2^, then developed using an AZ400k 1:4 developer (MicroChemicals), cleaned with water, rinsed, and dried by N_2_ gas flow. A 10 nm Ti adhesion layer, 100 nm Au layer, and 40 nm Pt layer were evaporated on the wafer using an electron-beam evaporator (Plassys MEB550S), and then the metal was lifted off in acetone to define the metal electrodes and interconnections. A top insulation layer of SU-8 2015 was then spin-coated at 5000 rpm for 1 min, soft based at 65°C for 3 min and 95°C for 5 min, and photolithography patterned, using MLA with a dose of 350 mJ/cm^2^ to expose the connection pads and to define the top insulation layer. After post-baking and a development procedure with the SU-8 developer, the wafer was cleaned with isopropanol and DI water, hard baked at 200°C, 180°C, and 150°C for 5 min each, and allowed to cool down below 95°C. The MEAs were lifted off from the wafer using a buffered oxide etchant (1:7) in an acid hood for about 4 h. An anchor hole was also patterned at the shank tip to facilitate the insertion of a 50 µm tungsten shuttle, to enable the handling and penetration of the flexible device into the spinal cord. Figure 1B shows the schematic of the flexible MEA fabrication.

### Glutamate sensor preparation

Fabricated electrodes were first characterized by electrochemical impedance spectroscopy (EIS) and cyclic voltammetry (CV) using an Autolab potentiostat (Metrohm, Herisau, Switzerland) with Nova 2.1.4 software to determine baseline functionality. EIS was measured from 10^5^ to 1 Hz, with 10 frequencies recorded per decade using a sine wave oscillation and a 0.01 V RMS voltage. CVs were scanned between -0.6 V and 0.8 V vs. an Ag/AgCl reference at 0.1 V/s.

Glutamate electrodes were prepared as described previously (19). To prepare the sensors, the Pt electrode sites were first cleaned by immersing them in 0.05 M H_2_SO_4_ (Fisher Scientific, Pittsburgh, PA, USA) and scanning from -0.35 V to 1.5 V at 1 V/s for 100 cycles. A screening layer of m-phenylenediamine (mPD, Sigma-Aldrich) was electrodeposited onto the electrode sites by immersing the electrodes in a 10 mM solution of mPD in 1x phosphate buffered saline (PBS) and applying 0.7 V for 20 minutes. Then, 1 µL of an enzyme solution containing 1 U glutamate oxidase (Cosmo Bio USA, Carlsbad, CA, USA), 13.7 mg bovine serum albumin (Fisher Scientific), 6.7 µL glutaraldehyde (Sigma-Aldrich) in 1 mL DI water was drop-casted onto 3 of the 6 electrode sites. These sites became the glutamate-sensitive sensor sites. Figure 2A shows a schematic of the electrode layers. A control solution lacking glutamate oxidase was dropped onto the remaining 3 electrode sites; these sites became the sentinel control sites.

Glutamate was detected amperometrically at 0.7 V vs. Ag/AgCl as the difference between the glutamate sensor sites and the sentinel control sites using a CH Instruments 1430 multichannel potentiostat (CH Instruments, Austin, TX, USA). To evaluate sensor functionality and sensitivity, electrodes were calibrated before and after use in standard concentrations of glutamate. In vivo, three glutamate-sensitive sites and one sentinel site were monitored simultaneously. A calibration curve was constructed from the post-calibration standards and used to convert detected in vivo current into concentration. To prepare reference electrodes for in vivo use, a silver wire was coated with silver chloride by immersing it into a 3 M solution of KCl and applying 4 V for 3 min.

### Animal Preparation

The study protocol was approved by the University of Pittsburgh Institutional Animal Care and Use Committee (IACUC). All experiments were performed in compliance with the National Institution of Health *Guide for the Care and Use of Laboratory Animals*.

Four male and four female Yorkshire pigs (44±2 kg) were used to evaluate the glutamate sensing probe performance and investigate the glutamate release probe profile during myocardial ischemia and reperfusion injury.

Combination of Tiletamine–Zolazepam (4 mg/kg, intramuscular) and Xylazine (2 mg/kg, intramuscular) was used to sedate the animal. The animals were given inhaled isoflurane 3-5% via a nose cone. When the animal was completely sedated and anesthetized, the animal was intubated and mechanically ventilated. After intubation, the anesthesia was reduced to 2-4% of inhaled isoflurane. General anesthesia was maintained with the isoflurane until the end of the preparation and then was switched to alpha chloralose (50 mg/kg initial bolus followed by a 20 mg/kg/h continuous infusion). Alpha chloralose limits the impact of anesthesia on the cardiac autonomic nervous system and cardiac myocardial excitability (3, 4, 49, 52). All animals were completely anesthetized during the whole procedure and the anesthesia level was adjusted throughout the experiment by monitoring corneal reflex, jaw tone, and hemodynamic indices. The animal temperature was maintained using water heating pads (T/PUMP; Gaymar Industries, Orchard Park, NY).

Heart rate, blood pressure, end-tidal carbon dioxide, and blood oxygen level were monitored continuously during the procedure. Arterial blood gas was tested hourly, or more frequently to prevent acid-base disorders, and ventilation rate/volume adjustment, or administration of sodium bicarbonate was performed as needed.

Electrocardiogram, systemic blood pressure, and left ventricular pressure were recorded using the Prucka CardioLab recording system (GE Healthcare, USA) and Millar system (ADInstruments, USA).

Carotid and femoral arteries, as well as external jugular and femoral veins, were catheterized for blood pressure monitoring and drug administration, respectively.

To expose the spinal cord, laminectomy was performed in these animals in the prone position. Upon completion of the laminectomy, the animals were switched to the supine position to perform the sternotomy procedure. Following sternotomy, the sternum was cauterized to avoid bleeding. The pericardial sac was cut and secured to the sternum using sutures. Once the heart was exposed, a Prolene suture was placed around the left anterior descending artery after its second diagonal branch. A nylon mesh epicardial sock electrodes were placed around the heart and a Millar catheter (Millar, USA) was placed in the left ventricle through the carotid artery.

The animals were placed in the lateral position to have access to both the heart and spinal cord to perform the myocardial ischemia intervention and the glutamate sensing probe insertion and recording.

At the end of the data collection protocol, animals were euthanized by inducing ventricular fibrillation via injection of potassium chloride under deep anesthesia.

### Experimental Protocols

Glutamate was detected in vivo using a 3-electrode setup. The reference electrode was a silver wire coated with silver chloride inserted into the muscle of the pig near the working electrode insertion site (53). A metal retractor was utilized as a counter electrode. Following 30 mins stabilizing period after the glutamate probe insertion, myocardial ischemia was performed for one hour by ligating the suture that was placed around the LAD. Two animals experienced ventricular arrhythmias during the 1-hour myocardial ischemia and the protocol could not be completed in these two animals. The suture was released after the 1-hour myocardial ischemia. Heart rate, systemic blood pressure, left ventricular systolic pressure, left ventricular contractility, activation recovery interval, dispersion of the repolarization, and glutamate concentration were recorded during the baseline, myocardial ischemia, and reperfusion.

### Hemodynamics Assessment

The lead II ECG was used to assess the heart rate during the protocol. Blood pressure was measured using the pressure transducer that was connected to the femoral arterial sheath. Left ventricular pressure was measured via a 5 French SPR-350 Millar Mikro-Tip pressure transducer catheter (Millar Instruments, Houston, TX) was inserted into the left ventricle and connected to an MPVS Ultra Pressure-Volume Loop System (Millar Instruments, Houston, TX). Left ventricular end-systolic pressure as well as left ventricular contractility (dP/dt max) were calculated using the left ventricular pressure signal. The hemodynamics parameters were analyzed using the Spike2 program (Cambridge Electronic Design) and (LabChart software; ADInstruments, Colorado Springs, CO)

### Activation Recovery Interval Analysis

A 56-electrode nylon mesh sock electrode was made using 0.2 mm silver wires and it was placed around the heart to measure the epicardial unipolar electrograms using a Prucka CardioLab electrophysiology mapping system (GE Healthcare, Fairfield, CT) (Figure 4D). Unipolar electrograms were filtered (0.05-500Hz) with the GE CardioLab System (54). Activation and repolarization time were detected using customized software (iScalDyn, University of Utah, Salt Lake City, UT), and the activation recovery intervals which is the difference between the repolarization and activation times were calculated (55). The activation recovery interval is a surrogate of local action potential duration (56, 57). Excessive sympathetic outflow shortens the ARIs and action potential duration.

### Dispersion of Repolarization

Dispersion of repolarization is defined as the variance of ARI among all 56 leads. Greater dispersion is associated with higher arrhythmia risk and therefore higher mortality.

### Arrhythmia score

We have calculated the arrhythmia score during the baseline, myocardial ischemia, and reperfusion to evaluate the arrhythmogenicity during ischemia-reperfusion injury. The arrhythmia score was calculated based on the following formula:(58) 0: <10 Premature ventricular contractions (PVCs), 1: 10–50 PVCs or 1 episode of VT or both, 2: >50 PVCs or >1 episode of VT or both, 3: SVF or 1 episode of NVF, 4: 2–4 episodes of NVF, 5: >4 episodes of NVF.

SVT (ventricular tachycardia) and SVF (ventricular fibrillation) denote the ventricular arrhythmias that terminated spontaneously and NVT (ventricular tachycardia) and NVF (ventricular fibrillation) denote the ventricular arrhythmias that did not terminate spontaneously.

### Statistical Analysis

For the normal paired data, paired t-test was performed. When the data did not pass the Shapiro-Wilk normality test, the Wilcoxon test was performed. All figures were created using GraphPad Prism software (version 8, GraphPad Software Inc, San Diego, CA) or MATLAB (2019 version, MathWorks). A P value ≤ 0.05 was considered statistically significant. Data are presented as median (25th, 75th percentiles) or mean ± SEM.

### Study Approval

The study protocol was approved by the University of Pittsburgh Institutional Animal Care and Use Committee (IACUC). All experiments were performed in compliance with the National Institution of Health *Guide for the Care and Use of Laboratory Animals*.

## Author contributions

S.S., E.R., X.T.C, and A.M. conceived and designed research; E.C fabricated the flexible probe. S.S, Y.K, E.R, and E.C conducted animal experiments; S.S, E.R, and Y.K. analyzed data. S.S and E.R drafted the manuscript and prepared figures; S.S, E.R, Y.K, E.C, X.T.C, and A.M interpreted the results of the experiments; All authors reviewed and approved the final version of the manuscript. The first authorship order position was listed based on intellectual contribution to the design of the study, performing the study, and interpretation of data.

## Acknowledgments

The study is funded by NIH R01 HL136836, NIH R01 NS110564, and NIH R21 DA049592. Siamak Salavatian is supported by the Competitive Medical Research Fund by University of Pittsburgh.

Elaine Robbins is supported by National Institute of Neurological Disorders and Stroke T32 NS086749 Training Grant

Xinyan Tracy Cui is supported by NIH R01 R01NS110564, R21DA049592

Aman Mahajan is supported by NIH R01 HL136836 and NIH R44 DA049630.

We thank Madison Fisher, Adrian Zalewski, and May Yoon Pwint for their technical assistance.

## REFERENCE

1. Myerburg RJ, and Junttila MJ. Sudden cardiac death caused by coronary heart disease. Circulation. 2012;125(8):1043–52.

2. Armour JA. Myocardial ischaemia and the cardiac nervous system. Cardiovascular research. 1999;41(1):41–54.

3. Dale EA, Kipke J, Kubo Y, Sunshine MD, Castro PA, Ardell JL, et al. Spinal cord neural network interactions: implications for sympathetic control of the porcine heart. Am J Physiol Heart Circ Physiol. 2020;318(4):H830–H9.

4. Omura Y, Kipke JP, Salavatian S, Afyouni AS, Wooten C, Herkenham RF, et al. Spinal Anesthesia Reduces Myocardial Ischemia-triggered Ventricular Arrhythmias by Suppressing Spinal Cord Neuronal Network Interactions in Pigs. Anesthesiology. 2021;134(3):405–20.

5. Exner DV, Pinski SL, Wyse DG, Renfroe EG, Follmann D, Gold M, et al. Electrical storm presages nonsudden death: the antiarrhythmics versus implantable defibrillators (AVID) trial. Circulation. 2001;103(16):2066–71.

6. Vaseghi M, Gima J, Kanaan C, Ajijola OA, Marmureanu A, Mahajan A, et al. Cardiac sympathetic denervation in patients with refractory ventricular arrhythmias or electrical storm: intermediate and long-term follow-up. Heart Rhythm. 2014;11(3):360–6.

7. Vaseghi M, and Shivkumar K. The role of the autonomic nervous system in sudden cardiac death. Prog Cardiovasc Dis. 2008;50(6):404–19.

8. Shen MJ, and Zipes DP. Role of the autonomic nervous system in modulating cardiac arrhythmias. Circ Res. 2014;114(6):1004–21.

9. Fukuda K, Kanazawa H, Aizawa Y, Ardell JL, and Shivkumar K. Cardiac innervation and sudden cardiac death. Circ Res. 2015;116(12):2005–19.

10. Armour JA. Cardiac neuronal hierarchy in health and disease. Am J Physiol Regul Integr Comp Physiol. 2004;287(2):R262–71.

11. Salavatian S, Ardell SM, Hammer M, Gibbons D, Armour JA, and Ardell JL. Thoracic spinal cord neuromodulation obtunds dorsal root ganglion afferent neuronal transduction of the ischemic ventricle. Am J Physiol Heart Circ Physiol. 2019;317(5):H1134–H41.

12. Cascio WE, Yang H, Muller-Borer BJ, and Johnson TA. Ischemia-induced arrhythmia: the role of connexins, gap junctions, and attendant changes in impulse propagation. J Electrocardiol. 2005;38(4 Suppl):55-9.

13. Mayer ML, and Westbrook GL. The physiology of excitatory amino acids in the vertebrate central nervous system. Prog Neurobiol. 1987;28(3):197–276.

14. Yung KK. Localization of glutamate receptors in dorsal horn of rat spinal cord. Neuroreport. 1998;9(7):1639–44.

15. Inquimbert P, Bartels K, Babaniyi OB, Barrett LB, Tegeder I, and Scholz J. Peripheral nerve injury produces a sustained shift in the balance between glutamate release and uptake in the dorsal horn of the spinal cord. Pain. 2012;153(12):2422–31.

16. Danbolt NC. Glutamate uptake. Prog Neurobiol. 2001;65(1):1–105.

17. Saether OD, Backstrom T, Aadahl P, Myhre HO, Norgren L, and Ungerstedt U. Microdialysis of the spinal cord during thoracic aortic cross-claming in a porcine model. Spinal Cord. 2000;28:153–7.

18. Watson CJ, Venton BJ, and Kennedy RT. In Vivo Measurements of Neurotransmitters by Microdialysis Sampling. Analytical Chemistry. 2006;78(5):1391–9.

19. Burmeister JJ, and Gerhardt GA. Self-Referencing Ceramic-Based Multisite Microelectrodes for the Detection and Elimination of Interferences from the Measurement of L-Glutamate and Other Analytes. Analytical Chemistry. 2001;73:1037–42.

20. Luan L, Wei X, Zhao Z, Siegel JJ, Potnis O, Tuppen CA, et al. Ultraflexible nanoelectronic probes form reliable, glial scar-free neural integration. Science Advances. 2017;3:1–9.

21. Rexed B. The cytoarchitectonic organization of the spinal cord in the cat. J Comp Neurol. 1952;96(3):414–95.

22. Howard-Quijano K, Yamaguchi T, Gao F, Kuwabara Y, Puig S, Lundquist E, et al. Spinal Cord Stimulation Reduces Ventricular Arrhythmias by Attenuating Reactive Gliosis and Activation of Spinal Interneurons. JACC Clin Electrophysiol. 2021;7(10):1211–25.

23. Weltman A, Yoo J, and Meng E. Flexible, penetrating brain probes enabled by advances in polymer microfabrication. Micromachines. 2016;7(10):180.

24. Nemani KV, Moodie KL, Brennick JB, Su A, and Gimi B. In vitro and in vivo evaluation of SU-8 biocompatibility. Materials Science and Engineering: C. 2013;33(7):4453–9.

25. He F, Lycke R, Ganji M, Xie C, and Luan L. Ultraflexible neural electrodes for long-lasting intracortical recording. Iscience. 2020:101387.

26. Zhao Z, Li X, He F, Wei X, Lin S, and Xie C. Parallel, minimally-invasive implantation of ultra-flexible neural electrode arrays. Journal of neural engineering. 2019;16(3):035001.

27. Castagnola E, Robbins EM, Wu B, Pwint MY, Garg R, Cohen-Karni T, et al. Flexible Glassy Carbon Multielectrode Array for In Vivo Multisite Detection of Tonic and Phasic Dopamine Concentrations. Biosensors. 2022;12(7).

28. Du ZJ, Kolarcik CL, Kozai TDY, Luebben SD, Sapp SA, Zheng XS, et al. Ultrasoft microwire neural electrodes improve chronic tissue integration. Acta Biomater. 2017;53:46–58.

29. Luan L, Robinson JT, Aazhang B, Chi T, Yang K, Li X, et al. Recent Advances in Electrical Neural Interface Engineering: Minimal Invasiveness, Longevity, and Scalability. Neuron. 2020;108(2):302–21.

30. Lotfi Marchoubeh M, Cobb SJ, Abrego Tello M, Hu M, Jaquins-Gerstl A, Robbins EM, et al. Miniaturized probe on polymer SU-8 with array of individually addressable microelectrodes for electrochemical analysis in neural and other biological tissues. Anal Bioanal Chem. 2021;413(27):6777–91.

31. Geramifard N, Dousti B, Nguyen CK, Abbott JR, Cogan S, and Varner V. Insertion mechanics of amorphous SiC ultra-micro scale neural probes. J Neural Eng. 2022.

32. Sharafkhani N, Kouzani A, D. Adams S, M. Long J, and O. Orwa J. A Pneumatic-based Mechanism for Inserting a Flexible Microprobe into the Brain. Journal of Applied Mechanics. 2022:1–20.

33. Singh S, Lo MC, Damodaran VB, Kaplan HM, Kohn J, Zahn JD, et al. Modeling the Insertion Mechanics of Flexible Neural Probes Coated with Sacrificial Polymers for Optimizing Probe Design. Sensors (Basel). 2016;16(3).

34. Kozai TD, Jaquins-Gerstl AS, Vazquez AL, Michael AC, and Cui XT. Brain tissue responses to neural implants impact signal sensitivity and intervention strategies. ACS Chem Neurosci. 2015;6(1):48–67.

35. Na K, Sperry ZJ, Lu J, Voroslakos M, Parizi SS, Bruns TM, et al. Novel diamond shuttle to deliver flexible neural probe with reduced tissue compression. Microsyst Nanoeng. 2020;6:37.

36. Stieglitz T, Beutel H, and Meyer J-U. A flexible, light-weight multichannel sieve electrode with integrated cables for interfacing regenerating peripheral nerves. Sensors and Actuators A: Physical. 1997;60(1-3):240–3.

37. O’Brien DP, Nichols TR, and Allen MG. Technical Digest MEMS 2001 14th IEEE International Conference on Micro Electro Mechanical Systems (Cat No 01CH37090). IEEE; 2001:216–9.

38. Wester BA, Lee RH, and LaPlaca MC. Development and characterization of in vivo flexible electrodes compatible with large tissue displacements. J Neural Eng. 2009;6(2):024002.

39. Moshayedi P, Ng G, Kwok JC, Yeo GS, Bryant CE, Fawcett JW, et al. The relationship between glial cell mechanosensitivity and foreign body reactions in the central nervous system. Biomaterials. 2014;35(13):3919–25.

40. Nguyen TNH, Nolan JK, Park H, Lam S, Fattah M, Page JC, et al. Facile fabrication of flexible glutamate biosensor using direct writing of platinum nanoparticle-based nanocomposite ink. Biosens Bioelectron. 2019;131:257–66.

41. Cao H, Li A-L, Nguyen CM, Peng Y-B, and Chiao J-C. An Integrated Flexible Implantable Micro-Probe for Sensing Neurotransmitters. IEEE Sensors Journal. 2012;12(5):1618–24.

42. Weltin A, Kieninger J, Enderle B, Gellner AK, Fritsch B, and Urban GA. Polymer-based, flexible glutamate and lactate microsensors for in vivo applications. Biosens Bioelectron. 2014;61:192–9.

43. Weltin A, Enderle B, Kieninger J, and Urban GA. Multiparametric, Flexible Microsensor Platform for Metabolic Monitoring In Vivo. IEEE Sensors Journal. 2014;14(10):3345–51.

44. Wang B, Wen X, Cao Y, Huang S, Lam HA, Liu TL, et al. An implantable multifunctional neural microprobe for simultaneous multi-analyte sensing and chemical delivery. Lab Chip. 2020;20(8):1390–7.

45. Schwerdt HN, Zhang E, Kim MJ, Yoshida T, Stanwicks L, Amemori S, et al. Cellular-scale probes enable stable chronic subsecond monitoring of dopamine neurochemicals in a rodent model. Commun Biol. 2018;1:144.

46. Zhao Z, Li X, He F, Wei X, Lin S, and Xie C. Parallel, minimally-invasive implantation of ultra-flexible neural electrode arrays. J Neural Eng. 2019;16(3):035001.

47. Howard-Quijano K, Takamiya T, Dale EA, Kipke J, Kubo Y, Grogan T, et al. Spinal cord stimulation reduces ventricular arrhythmias during acute ischemia by attenuation of regional myocardial excitability. Am J Physiol Heart Circ Physiol. 2017;313(2):H421–H31.

48. Vaseghi M, Yamakawa K, Sinha A, So EL, Zhou W, Ajijola OA, et al. Modulation of regional dispersion of repolarization and T-peak to T-end interval by the right and left stellate ganglia. American Journal of Physiology-Heart and Circulatory Physiology. 2013;305(7):H1020–H30.

49. Howard-Quijano K, Takamiya T, Dale EA, Yamakawa K, Zhou W, Buckley U, et al. Effect of Thoracic Epidural Anesthesia on Ventricular Excitability in a Porcine Model. Anesthesiology. 2017;126(6):1096–106.

50. Vaseghi M, Lux RL, Mahajan A, and Shivkumar K. Sympathetic stimulation increases dispersion of repolarization in humans with myocardial infarction. American Journal of Physiology-Heart and Circulatory Physiology. 2012;302(9):H1838–H46.

51. Salavatian S, Beaumont E, Gibbons D, Hammer M, Hoover DB, Armour JA, et al. Thoracic spinal cord and cervical vagosympathetic neuromodulation obtund nodose sensory transduction of myocardial ischemia. Autonomic neuroscience : basic & clinical. 2017;208:57–65.

52. Batul SA, Olshansky B, Fisher JD, and Gopinathannair R. Recent advances in the management of ventricular tachyarrhythmias. F1000Res. 2017;6:1027.

53. Robbins EM, Castagnola E, and Cui XT. Accurate and stable chronic in vivo voltammetry enabled by a replaceable subcutaneous reference electrode. iScience. 2022;25(8):104845.

54. Vaseghi M, Yamakawa K, Sinha A, So EL, Zhou W, Ajijola OA, et al. Modulation of regional dispersion of repolarization and T-peak to T-end interval by the right and left stellate ganglia. Am J Physiol Heart Circ Physiol. 2013;305(7):H1020–30.

55. Ajijola OA, Vaseghi M, Zhou W, Yamakawa K, Benharash P, Hadaya J, et al. Functional differences between junctional and extrajunctional adrenergic receptor activation in mammalian ventricle. Am J Physiol Heart Circ Physiol. 2013;304(4):H579–88.

56. Millar CK, Kralios FA, and Lux RL. Correlation between refractory periods and activation-recovery intervals from electrograms: effects of rate and adrenergic interventions. Circulation. 1985;72(6):1372–9.

57. Chinushi M, Tagawa M, Kasai H, Washizuka T, Abe A, Furushima H, et al. Correlation Between the Effective Refractory Period and Activation-Recovery Interval Calculated From the Intracardiac Unipolar Electrogram of Humans With and Without dl-Sotalol Treatment. JAPANESE CIRCULATION JOURNAL. 2001;65(8):702–6.

58. Curtis MJ, and Walker MJ. Quantification of arrhythmias using scoring systems: an examination of seven scores in an in vivo model of regional myocardial ischaemia. Cardiovasc Res. 1988;22(9):656–65.

