## Supplemental figures for "Real-time in vivo thoracic spinal glutamate sensing reveals spinal hyperactivity during myocardial ischemia"

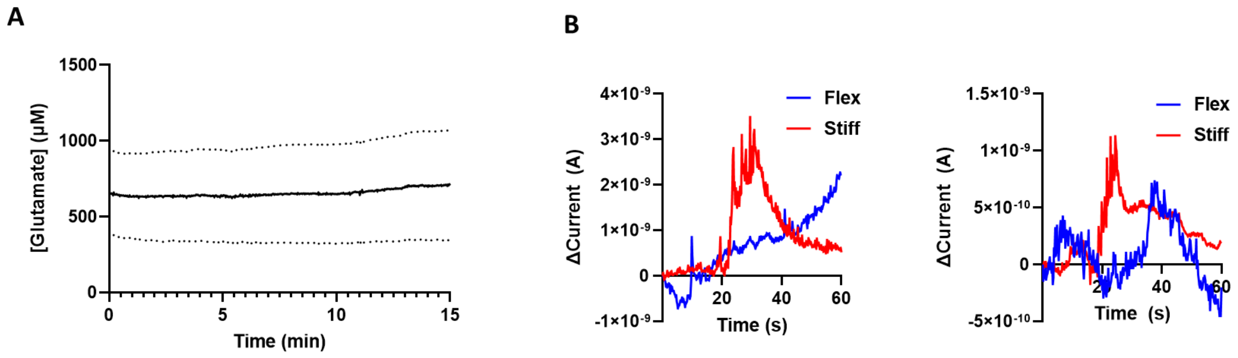


**Supplemental Figure 1. Glutamate electrode performance.** (**A**) The average of the recorded glutamate concentration using both sharpened shuttle and PEG-assisted probe (**B**) Two trials of data recorded during motion artifacts using a stiff acute Neuronexus probe and flexible probe, both inserted and recorded at the same time.

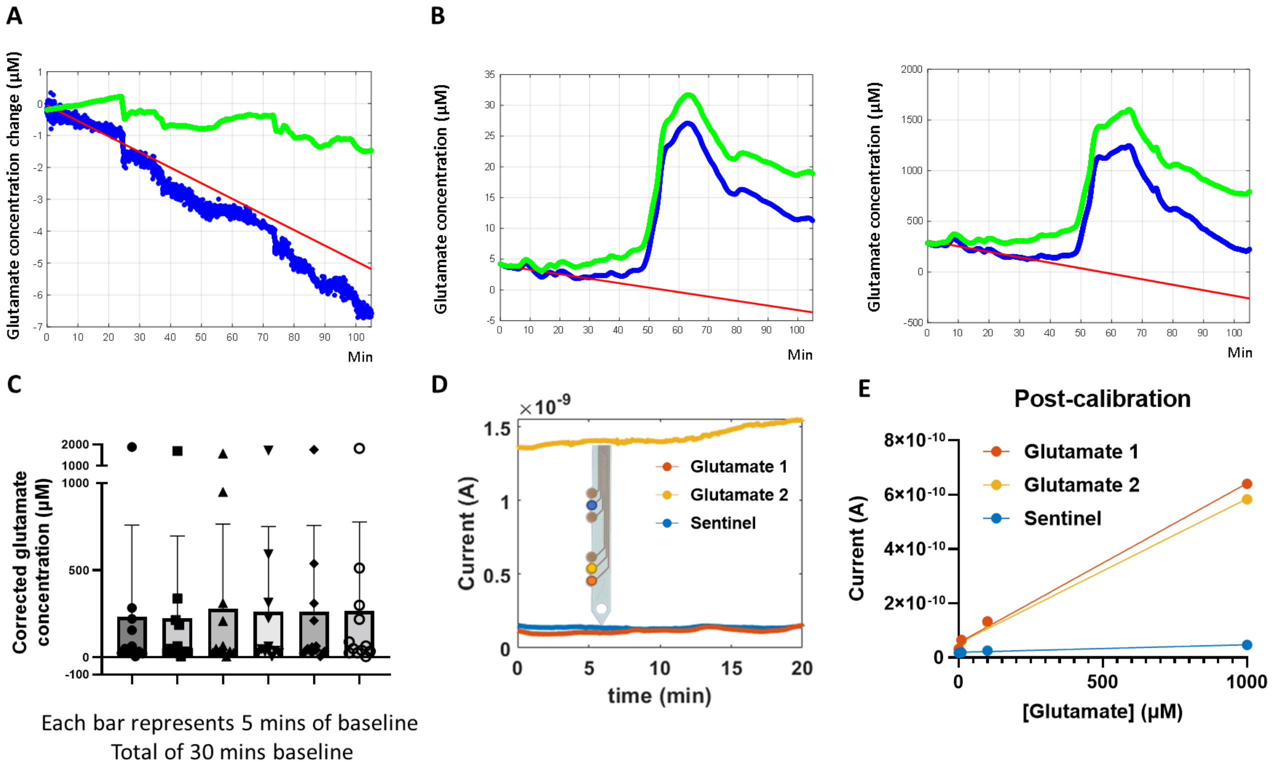


**Supplemental Figure 2. In vivo sensitivity of the glutamate probe** (**A**) change in the glutamate concentration from baseline at time=0 (blue trace). The Red line is fitted on the first 30 minutes of the data and its slope is subtracted from the data. The green trace shows the data after correcting it for the slight loss of sensitivity and it oscillates around 0 µM. **(B)** Two examples of correcting the data for the slight loss of sensitivity. Blue traces are representing the data without the loss of sensitivity correction while the green traces are the corrected data after removing the slope of the red fitted line. **(C)** Corrected glutamate concentration during the 30 minutes of baseline. No significant change is observed during the baseline (n=12 electrodes, P= 0.61, Friedman test). **(D)** Representative glutamate signaling from 3 different sites (2 glutamate sensing sites and a sentinel) **(E)** Post-experiment calibration of electrodes in D confirming the electrodes are functional.
